# Building bridges from genome to physiology using machine learning and Drosophila experimental evolution

**DOI:** 10.1101/2022.07.18.500543

**Authors:** James N. Kezos, Thomas T. Barter, Mark. A. Phillips, Larry G. Cabral, Grigor Azatian, José Buenrostro, Punjot Singh Bhangoo, Annie Khong, Gabriel T. Reyes, Adil Rahman, Laura A. Humphrey, Timothy J. Bradley, Laurence D. Mueller, Michael R. Rose

## Abstract

Drosophila experimental evolution, with its well-defined selection protocols, has long supplied useful genetic material for the analysis of functional physiology. While there is a long tradition of interpreting the effects of large-effect mutants physiologically, in the genomic era identifying and interpreting gene-to-phenotype relationships has been challenging, with many labs not resolving how physiological traits are affected by multiple genes throughout the genome. Drosophila experimental evolution has demonstrated that multiple phenotypes change due to the evolution of many loci across the genome, creating the scientific challenge of sifting out differentiated but noncausal loci for individual characters. The fused lasso additive model method (FLAM) allows us to infer some of the differentiated loci that have relatively greater causal effects on the differentiation of specific phenotypes.

The experimental material used in the present study comes from 50 populations that have been selected for different life-histories and levels of stress resistance. Differentiation of cardiac robustness, starvation resistance, desiccation resistance, lipid content, glycogen content, water content, and body masses was assayed among 40 to 50 of these experimentally-evolved populations. Through FLAM, we combined physiological analysis from eight parameters with whole-body pooled-seq genomic data to identify potentially causally linked genomic regions. We have identified approximately 1,900 significantly differentiated 50 kb genomic windows among our 50 populations, with 161 of those identified genomic regions highly likely to have a causal effect connecting specific genome sites to specific physiological characters.

## Introduction

The genomic era has provided biologists with powerful tools for surveying variation at the levels of DNA sequences, RNA transcripts, and even metabolites. While there is a long tradition of interpreting the effects of large-effect mutants physiologically, in the genomic era identifying and interpreting gene-to-phenotype relationships has been challenging. It is still the case that some phenotypes are interpretably associated with large-effect genetic mutations such as *opa1* (e.g. Shahrestani et al. 2009). But that research does not address the genetic foundations of physiological traits that are affected by multiple genes distributed throughout the genome. Methods such as GWAS were once seen as a potential solution to this challenge, but research with such analytical methods has failed to account for most of the heritable variation for functional characters. Fortunately, we now have a range of new tools that come from statistical learning research, which provide potential solutions to the challenges of genome-wide analysis of functional characters.

Drosophila experimental evolution has long supplied useful genetic material for the analysis of functional physiology (e.g. Burke 2010; Burke and Rose 2009; Shahrestani 2011). There are quite a few advantages to using model organisms subjected to sustained experimental evolution in the lab. Not only does experimental evolution provide well-defined replication and selection protocols, but it also usually yields well-documented evolutionary histories for the populations under study (Garland and Carter 1994; Garland and Rose 2009, Rose et al. 2004;. Perhaps most importantly, laboratory selection can produce stronger phenotypic differentiation than the phenotypic differences observed within or among natural populations (Garland and Rose 2009; Rose et al. 2004). The challenge with analyzing the physiological genetics of Drosophila experimental evolution is that multiple phenotypes change due to the evolution of many loci across the genome (Rose et al. 2004; Burke et al. 2010; Graves et al. 2017). Thus, determining the genetic foundations of a specific physiological phenotype is more complicated than simply looking for all genomic differentiation arising with a particular experimental evolution paradigm.

At its core, the scientific challenge is how to sift out differentiated but noncausal loci when inferring the genes that affect a particular phenotype, given well-replicated genome-wide differentiation. Fortunately, Mueller et al. (2018) used numerical simulations and pilot tests to demonstrate the potential of machine learning to address this challenge. The specific type of machine learning employed by Mueller et al. (2018) was the fused lasso additive model method (FLAM; Petersen et al. 2016). Mueller et al. (2018) showed that it is possible to infer at least some of the differentiated loci causally associated with observed phenotypic response(s) to experimental evolution, while rarely mistakenly identifying non-causal variants.

The experimental material that we use in the present study comes from populations that have been selected for different life-histories and stress resistance (Rose et al. 2004; Burke et al. 2016; Kezos et al. 2019). It is important to note that, though populations with different life-cycle timings are not specifically selected for different levels of stress resistance, they do nonetheless show distinctive enhancements in stress and performance characters. For example, early on it was found that a variety of stress resistances and performance characters were improved in experimentally evolved stocks with postponed senescence (vid. Service et al.Luckinbill et al. 1984; Rose et al. 1984; Luckinbill and Clare 1985; Service et al. 1985; Luckinbill et al. 1988; 1988; Graves and Rose 1990; Graves et al. 1992; Rose et al. 2004; Swallow et al. 2009). Resistance to starvation, desiccation, and pacing-induced cardiac arrest, along with increased metabolic components (i.e. lipid, glycogen, water content) and flight endurance are all associated with selection for postponed senescence (Service et al. 1985, Graves et al. (1988) as well as Graves and Rose 1990, Graves et al. 1992; Gibbs et al. 1997; Chippindale et al. 1997; Djawdan et al. 1998, Gibbs and Gefen 2009; Kezos et al 2017, Kezos et al. 2019). Further work on the evolutionary physiology of these *Drosophila* has revealed the marked differences in physiological machinery underlying such independently evolving functional characters. Adult starvation resistance, for example, depends overwhelmingly on total stored calories (e.g. Djawdan et al. 1998), while desiccation resistance depends chiefly on water content and rate of water loss (e.g. Gibbs et al. 1997; Gibbs and Gefen 2009). These two characters underpinning desiccation resistance were further shown to evolve separately from each other when sustained strong selection for desiccation resistance is applied (Archer et al. 2007). Overall, we would say that this is one of better established laboratory systems for the study of evolutionary physiology.

Through our labs’ machine learning approach (FLAM), we have combined physiological analysis with whole-body pooled-seq genomic data (vid. Burke et al., 2010) to identify genomic regions which might be causally linked to specific physiological phenotypes. That is, we have identified specific genomic regions that are likely to have a causal effect connecting genome to physiology.

## Materials and Methods

### 3.1 Populations Used

This study employed 50 of the large, outbred, and highly differentiated populations created by the Rose laboratory since 1980 (e.g. Rose 1984; Rose et al. 1992; Rose et al. 2004). All of the populations assayed here descend from a single *Drosophila melanogaster* population, called IV. The IV population originated in 1975 as a sample of *D. melanogaster* caught in Amherst, Massachusetts. After four and a half years of laboratory culture, the B_1-5_ populations (baseline) and O_1-5_ populations (70-day generation cycle) were derived from the single IV population in 1980 (Rose 1984; Rose et al. 2004). The 50 populations studied here are referred to as ACO_1-5_, AO_1-5_, B_1-5_, BO_1-5_, TDO_1-5_, TSO_1-5_, CO_1-5_, nCO_1-5_, SCO_1-5A_, and SCO_1-5B_. These large, outbred populations were maintained on a banana medium (agarose, banana, light and dark corn syrup, barley malt, yeast, ethanol and water). with a 24L:0D light cycle. All populations were kept at moderately large census populations sizes (N > 1,000) to avoid confounding inbreeding effects.

The ACO_1-5_ populations, for which “A” stands for accelerated and has a generation cycle length of 10 days, were derived from the CO_1-5_ populations in 1991 (Chippendale et al 1997; Chippendale et al 2004). The CO_1-5_ populations, which have a generation cycle length of 28 days, were derived from the O_1-5_ populations in 1989 (Rose et al 1992). The AO_1-5_, BO_1-5_, and NCO_1-5_ populations were all derived from the O_1-5_ populations in the years 2008, 2007, and 2009, respectively (Burke et al 2016; Graves et al. 2017). These populations are the newly-derived versions of the ACO, B, and CO populations. Each new population was founded from the same number replicate of their ancestral population. The ACO and AO populations fall under the A-type selection regime, whereas the B and BO populations fall under the B-type selection regime, and lastly, the CO and NCO populations fall under the C-type selection regime.

In 1988, two sets of populations were derived from the O_1–5_ populations. One set (D_1–5_) was selected for desiccation resistance while the other set (C_1–5_) was maintained to control for desiccation resistance selection (Rose et al., 1992; Phillips et al 2018). The C_1–5_ populations were handled like the D_1–5_ populations, except flies were given nonnutritive agar instead of desiccant [18]. In 2005, these populations were relaxed from selection and kept on a 21-day culture regime to the present day. Under this new regime, the D populations have been renamed to TDO, and the C populations to TSO. In total, the TDO populations underwent ∼ 260 generations of selection for desiccation. These ten populations fall under the T-type selection regime.

The SCO_1-5A_ and SCO_1-5B_ populations (28-day generation cycle) are ten populations intensely selected for starvation resistance. The ten SCO populations were derived from the five CO populations in August 2010. From each CO population, two SCO populations were derived. The S-type selection regime and generation cycle is described in detail in Kezos et al. 2019. In short, each population is exposed to nonnutritive agar until a 75-80% mortality threshold has been reached. When the 75% mortality threshold is achieved, the surviving adult flies serve as founders for the next generation. At the beginning of this experiment, the starvation period took just three days to achieve 75-80% mortality. Currently, the starvation period lasts for approximately 10 days. The group of populations subjected to this selection regime is referred to as the S-type treatment group.

### 3.2 Functional Assay Methods

#### Rearing protocols

Two run-in generations of 14-day life-cycles were used to remove any parental or grand-parental epigenetic effects not related to the selection regime that could potentially affect the measured physiological responses. The populations were cultured in banana medium from egg to adult, on a 24L:0D light schedule. A petri dish with the banana medium was placed into each respective cage. Adult flies were given 24 hours to lay eggs on the media. From the plate, eggs were collected at a density of 60 to 80 eggs and subsequently placed in a vial containing fresh media. At the end of each run-in generation (day 14 from egg), the populations were transferred to an acrylic cage. Replicate populations of the same number were handled in parallel at all stages. On day 14 of the second run-in generation, the adults were assigned at random to one of the following eight assays.

At the beginning of the 14-day from egg desiccation resistance and starvation resistance assays, the CO and nCO populations had been under 325 and 72 generations of C-type selection. The SCO_A_ and SCO_B_ populations had completed 55 generations of intense selection. For the 14-day from egg cardiac arrest rate assay, the CO and nCO populations were under selection for 332 and 79 generations, respectively. The SCO_A_ and SCO_B_ populations had completed 62 generations of intense selection. At the beginning of this mortality assay, the CO and nCO populations had been under C-type selection for 341 and 88 generations. The SCO_A_ and SCO_B_ populations had completed 70 generations of intense selection. For the 14-day from egg glycogen content, lipid content, and water content assays, the CO and nCO populations had been under 352 and 98 generations of selection. The SCO_A_ and SCO_B_ populations had completed 81 generations of intense selection. For the age-specific cardiac arrest rate and lipid content assays, the CO and SCO_A_ populations had been under their respective selection regime for 357 and 86 generations, respectively.

#### Glycogen Content

For each population, six groups of 10 females (14 days old from egg) were anesthetized using ethyl ether, placed in 1.7 milliliter microcentrifuge tubes, and stored in a freezer over night. The next day, each group was placed in aluminum weighing boats and placed in an oven for one hour at 60 °C. Each group was transferred to their respective microcentrifuge tube where 700 microliters of deionized water was added to each tube. The flies were then ground using a hand-held battery-operated grinder. Each tube was boiled in water for five minutes. Once complete, 500 microliters of the supernatant was carefully pipetted into a new microcentrifuge tube to minimize noise from precipitate contamination. Each tube was then vortexed to thoroughly mix the contents, then 100 microliters from each tube was transferred to a 13 x 100- millimeter test tube. Three milliliters of an anthrone reagent was added to each test tube. The anthrone reagent composition was 150 milligrams of anthrone per 100 milliliters of 72% sulfuric acid. Each test tube was then incubated in a water bath set to 90 °C for 10 minutes. Two one-milliliter samples from each test tube were placed in their own cuvettes. Absorbance was then measured using a Perkin Elmer Lambda spectrophotometer at a wavelength of 620 nanometers. All measurements were taken within 10 minutes of being removed from the water bath. Five controls of known glycogen concentration underwent the same process.

#### Water Content

For each population, six groups of 10 females (14 days old from egg) were anesthetized using ethyl ether. Each group was placed in aluminum weighing boats, and had their wet mass measured. The flies were frozen overnight and placed in a drying oven the next day at 60 °C for 24 hours. Each group was then reweighed. The difference between the wet mass and dry mass represented the water content of that group.

#### Lipid Content

For each population, lipid content was measured for six groups of 10 females at age 14 days from egg. After the dry masses of the groups were recorded for the water content assay, each group was placed in their own Whatman thimble. The thimbles were placed in the extractor of a Soxhlet apparatus. Petroleum ether was used as the extraction solvent. The thimbles were in the Soxhlet apparatus for 24 hours, and upon completion, were removed and placed in a drying oven at 60 °C for one hour. The post-extraction mass of each group was recorded using a microbalance scale. The difference between the dry mass and the post-extraction mass of each group represented the specific group’s lipid content.

A second set of lipid content measurements were made for the five CO and five SCO_A_ populations at six different adult ages. These measurements were conducted at ages 14, 21, 28, 35, 42, and 49 days from egg. The protocol is the same as the previous paragraph.

#### Desiccation resistance assay

Thirty individual female flies (14 days old from egg) from each population were placed in their own desiccant straw. A piece of cheesecloth separated the fly from the pipet tip at the end of the straw that contained 0.75 grams of desiccant (anhydrous calcium sulfate). The pipette tip containing desiccant was sealed with a layer of Parafilm. Mortality was checked hourly, using lack of movement under provocation as a sign of death. Note that this was a materially different procedure than the one we have employed previously in our studies of desiccation resistance (e.g. Service et al. 1985; Graves et al. 1992; Djawdan et al. 1998), which used vials.

#### Starvation resistance assay

Thirty individual female flies (14 days old from egg) from each population were placed in their own starvation straw with agar. The agar plug provides adequate humidity, but no nutrients. Mortality was checked every four hours, using lack of movement under provocation as a sign of death.

#### Cardiac Arrest Rate Assay

Forty female flies (14 days old from egg) from each replicate per stock were chosen at random. The flies were anesthetized for three minutes using triethylamine, also known as FlyNap, and then placed on a microscope slide prepared with foil and two electrodes. FlyNap was chosen as the anesthetic because of its minimal effect on heart function and heart physiology when administered for more than one minute, but less than the lethal time of five minutes (Chen and Hillyer 2013). The cold-shock method was not used as an anesthetic for the cardiac pacing assay, because the flies need to be fully anesthetized throughout the procedure. If the flies regain consciousness, the added stress and abdominal contractions while trying to escape would alter heart rate and function more than the side-effects of FlyNap’s do. Paternostro et al. (2001) found that FlyNap is less disruptive to cardiac function than the two other substances commonly used for *Drosophila* anesthesia, carbon dioxide and ether. Two electrodes were attached to a square-wave stimulator in order to produce electric pacing of heart contraction. Anesthetized flies were attached to the slide between the foil gaps using a conductive electrode jelly touching the two ends of the fly body, specifically the head and the posterior abdomen tip. The electrical shock settings for this assay were 40 volts, six Hertz, and 10 ms pulse duration. The settings of the electrical pacing assay were chosen in order to increase the contraction rate to such a high enough level that cardiac arrest would be consistently induced. Using an inverted compound microscope, an initial check of whether the heart was contracting or in arrest was made immediately after completion of the 30 second shock. A second check was made after a two-minute “recovery” period to determine if the heart was in cardiac arrest. The protocol for this assay was originally outlined in Wessells and Bodmer (2004).

A second set of cardiac arrest rate measurements were made for the CO and SCO_A_ populations at six different adult ages. These measurements were conducted at ages 14, 21, 28, 35, 42, and 49 days from egg. The protocol is the same as the above-mentioned 14-day from egg pacing experiment.

### 3.3 Statistical Methods

#### Analysis for Physiological Characters

Each of the five measured phenotypes, glycogen content, lipid content, water content, starvation resistance, and desiccation resistance, were analyzed separately. We now outline the two linear mixed-effects models we used for starvation resistance.

The first model was used for our within selection regime comparisons (CO vs nCO and SCO_A_ vs SCO_B_). Let *y_ijk_* be the measured starvation resistance for selection treatment-*i* (*i*=1 (CO or SCO_A_) and 2 (nCO or SCO_B_)), population-*j* (*j*= 1..10), and individual-*k* (*k*=1..*n_l_*). Then the effects of the fixed and random effects can be modeled as,

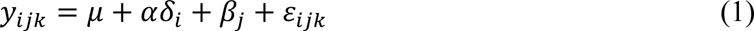

where 8*_i_*=0 if *i*=1 and 1 otherwise.

The second model was used for our selection type comparisons (i.e. C-type selection vs. S-type selection). Let *y_ijk_* be the measured starvation resistance for selection type-*i* (*i*=1 (C type) and 2 (S type)), population-*j* (*j*= 1..20), and individual-*k* (*k*=1..*n_l_*). Then the effects of the fixed and random effects can be modeled as,

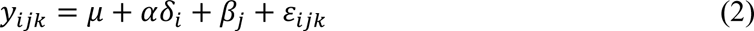

where 8*_i_*=0 if *i*=1 and 1 otherwise.

For these two models, the main effects of selection regime and replicate population were measured by α and 𝛽, respectively. The different populations contributed random effects to these measurements by genetically based differences that arise due to random genetic drift and were measured by 𝛽 while individual random variation was measured by ε. Both sources of random variation were assumed to be independent normally distributed random variables with zero means. The model parameters were estimated with the R *lme* function (R Core Team 2015).

Cochran-Mantel-Haenszel (CMH) tests were used to analyze the rates of cardiac arrests between two different stocks (i.e. CO vs nCO). The CMH test is used when there are repeated tests of independence, or multiple 2×2 tables of independence. This is the equation for the CMH test statistic, with the continuity correction included, that we used for our statistical analyses:

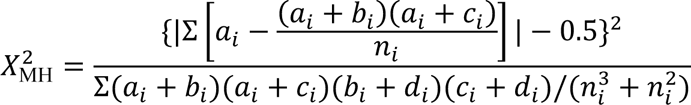

We designated “a” and “b” as the number of cardiac arrests in population *i* of the first stock and population *i* of the second stock, respectively. We designated “c” and “d” as the number of contracting hearts in the two populations. The n*_i_* represented the sum of a*_i_*, b*_i_*, c*_i_*, and d*_i_*. The subscript *i* (i = 1..5), represented one of the five replicate populations within each of the four stocks.

### 3.4 Pooled Genome Sequencing and Analysis

The goal of genomic analysis was to identify patterns of SNP differentiation between the 50 populations that predict the aforementioned physiological parameters. The genomic data utilized in this study have been previously analyzed and published, however, the present analysis linking our genomic analysis to the physiological differentiation is novel. SNP data came from previously published pool-seq DNA data from Graves et al., 2017 (A, B, and C populations), Kezos et al., 2019 (S populations), and Phillips et al., 2018 (T populations).

Genomic data was reprocessed since the original studies did not include all populations used for this study together at once. However, processing steps were otherwise the same. Briefly, fastq files corresponding to each population were mapped to the *D. melanogaster* reference genome (version 6.14) with BWA using bwa mem with default setting, and SAMtools was used to convert the resulting SAM files to BAM files, remove potential PCR duplicates, and merge all BAM files into a single mpileup (Li and Durbin 2009; Li et al 2009). PoPoolation2 was used to convert this mpileup file to a simplified file format that contains counts for all bases in the reference genome and for all populations being analyzed. (Kofler et al. 2011). RepeatMasker V4.12 (http://www.repeatmasker.org) and PoPoolation2 were then used to identify and remove highly repetitive genomic regions where proper read mapping is difficult. Lastly, SNPs were called based on the following criteria: minimum coverage of 20x and maximum of 200x in each population, and a combined minor allele frequency of 2% across all populations. This resulted in a SNP table with ∼781 K sites.

To link patterns of physiological and genomic differentiation, the generalized Cochran– Mantel–Haenszel (CMH) test (Landis et al 1978) was first used to identify SNPs that were significantly differentiated between the 50 populations (Rstudio; mantelhaentest function). Tests were performed for each SNP in the data set, followed by correction for multiple comparisons. To do this, the “plug in method” (Hastie et al., 2009) was used. Briefly, suppose we have M total hypothesis tests, and we let V be the number of false positives and S be the number of true positives, then the false discovery rate (FDR) is defined as V/(V + S). A critical test statistic, C, can be chosen and the plug-in method computes the FDR for that critical point. V + S is estimated to simply be the total number of test statistics from the M hypothesis tests that exceed C. V can in turn be estimated through permutation. Here, that was done by essentially shuffling population labels then performing the CMH test at each polymorphic site in the permuted data set. The number of significant test statistics greater than all values of C was then recorded. This was repeated 100 times to ensure accurate estimates of V for values of C. Once V and V + S were estimated, the FDR was calculated for each value of C. Using these results, we calibrated to a conservative FDR of 0.005 (i.e., significance threshold is the value of C that gives an FDR of 0.005).

### 3.5 Fused Lasso Additive Model Analysis (FLAM)

After identifying SNPs that were differentiated across selection regimes, a statistical learning approach called the “fused lasso additive model” or “FLAM” (Petersen et al 2016; Mueller et al 2018) was used to determine which of the genomic regions these SNPs represent best predict patterns of differentiation for our physiological parameters. Here, the assumption is that these genomic regions are the ones most likely to be casually linked to relevant patterns of physiological differentiation. Due to genetic linkage, it is not necessary to consider individual significant SNPs. Instead, a list of the most significant SNP for every 50 KB genomic window was generated and these markers were used in the FLAM analysis. This mimics the FLAM approach described in Mueller et al. (2018). As in Mueller et al. (2018), this is accounted for using a permutation procedure where each FLAM analysis is run multiple times and the order of potential predictor variables is randomly shuffled. The final list of “best predictors” consists of genetic loci that occur at commonly identified across permutations. In this study, a total of 100 permutations were run for each parameter, and the final list of best predictors consisted of loci that showed up in at least 50% of permutations of a given parameter.

After determining which genomic regions best predict observed physiological differences between populations, a list of genes associated with each region was generated. Given that predictors are markers representing genomic regions, it is not assumed that the SNP markers are themselves causal. Given that the SNPs used in the FLAM analyses are the most significant SNPs in their respective genomic region, we expect them to be relatively close to the true causative sites (Baldwin-Brown et al. 2014). Candidate genes were therefore defined as those in 2 KB windows around each SNP marker.

After generating a list of candidate genes associated with each parameter, publicly available online tools, Flybase (www.flybase.org) and g:Profiler (https://biit.cs.ut.ee/gprofiler/gost), were used to identify associated gene ontology (GO) terms and KEGG pathways. Batch downloads of gene profiles from Flybase were downloaded. With g:Profiler, the tool performs functional enrichment analysis using our identified genes. For each physiological parameter, a different query was performed with that specific trait’s respective set of genes. For comparative purposes, the genes were ran across 1) only annotated genes AND 2) all known genes, and a Benjamini-Hochberg FDR correction was used. GO terms and KEGG pathways with a significant enrichment (0.05 threshold) is found in Table S7.

## Results

### 4.1 Starvation Resistance and Lipid Content

Figure 1 and Table 1 display the mean starvation resistance of the 50 populations, respectively. As in previous studies (e.g. Djawdan et al. 1998), populations with a longer generation cycle were observed to have relatively increased starvation resistance (Fig. 1B). The A-type populations had the shortest starvation survival times and lowest lipid content (34.142 hours and 0.02 mg/fly), followed by the B-type populations (46.63 hours and 0.032 mg/fly), then the C-type populations (64.142 hours and 0.054 mg/fly), then the T-type populations (71.293 hours), and lastly, the S-type populations (151.671 hours and 0.14 mg/fly; Table S1). The five selection regimes all significantly differentiated from each other on a pairwise basis, as determined using post hoc EMmeans analysis, with the exception of C-type versus T-type comparison (p-values of 0.253 and for starvation resistance; Table S2). A-type versus B-type comparison was barely insignificant with a p-value of 0.056. In addition, when analyzing the pairwise differences between long-established and recently derived populations within the A-, B, and C-type selection regimes, there are no statistically significant differences for starvation resistance (e.g. the average ACO female starvation time is 34.094 hours while the AO average is 34.19 hours, p-value = 0.929; Table S3). The same is true for comparing the five TSO and five TDO populations against each other as well as the five SCO-a and five SCO-b populations. Similar comparisons were found regarding lipid content with the exception of the five ACO and five AO populations (p-value = 0.019). Because there is little to no statistical difference within the five selection regime, we can now treat each selection regime five 10-fold replicated sets of populations, thus increasing the statistical power of our experimental and genomic testing. In the T-type selection regime, there is no longer a difference in starvation resistance between the once intensely selected desiccation resistant TDO populations (73.537 survival hours) and their respective mildly selected starvation resistant TSO populations (69.094 survival hours, p-value = 0.152). Similar to the A, B, and C-type populations, we will refer to these 10 populations as the T-type populations.

**Figure 1.**
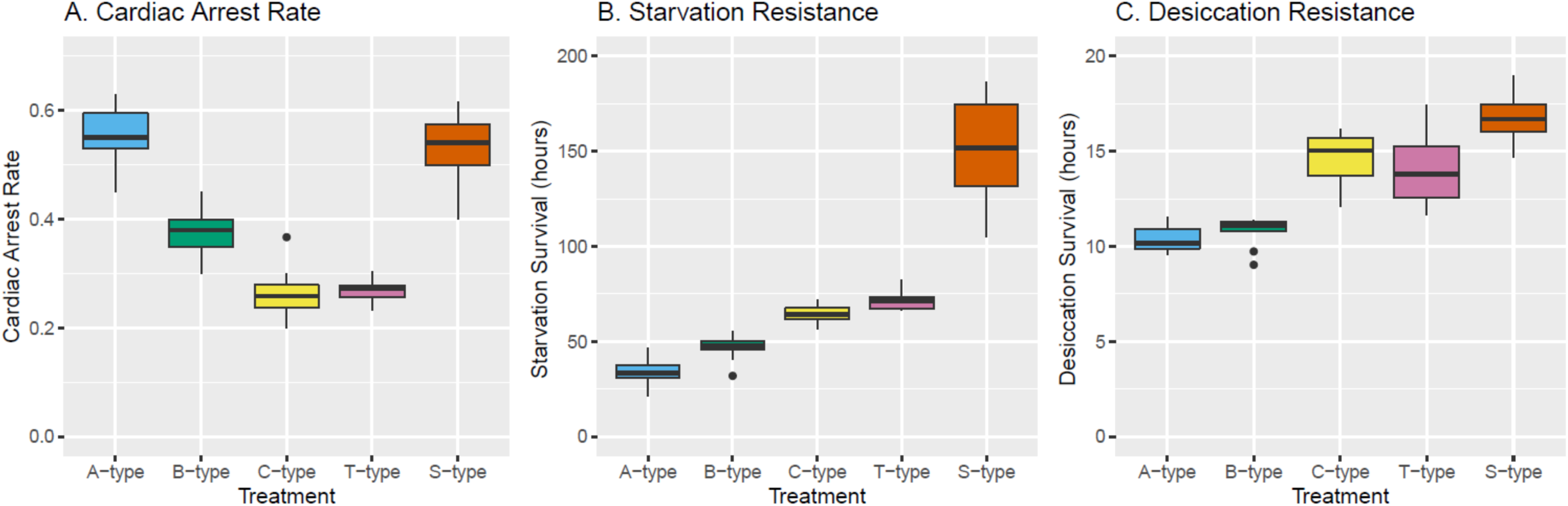
Summary boxplots of (A) cardiac arrest rates, (B) starvation resistance, and (C) desiccation resistance for the five selection regimes. Regarding cardiac arrest rates following electrical pacing, we see significant decline in rates as you increase generation cycle length. However, the S-type regime, which have increased starvation resistance and therefore, increased lipid content, have significantly higher arrest rates than B-, C-, and T-type regimes (Cochran-Mantel-Haenszel test). Regarding survival during starvation (B) and desiccation (C), we see an increase in survival times across populations with longer generation cycle lengths. The S-type populations have the longest survival times during starvation and desiccation (R-studio linear mixed effects model analysis).

**Table 1.** Phenotypic Characterization of C-type and S-type Populations

| | A-type Average<br>(mean ± SE) | B-type Average<br>(mean ± SE) | C-type Average<br>(mean ± SE) | S-type Average<br>(mean ± SE) | T-type Average<br>(mean ± SE) |
| --- | --- | --- | --- | --- | --- |
| Cardiac Arrest Rate (%) | 55.25 ± 1.766 | 37.5 ± 1.443 | 26.33 ± 3.5 | 53.33 ± 2.9 | 26.7 ± 0.721 |
| Survival during Starvation (h) | 34.142 ± 2.307 | 46.63 ± 2.03 | 64.14 ± 1.23 | 151.67 ± 4.23 | 71.293 ± 1.543 |
| Survival during Desiccation (h) | 10.383 ± 0.228 | 10.785 ± 0.248 | 14.67 ± 0.24 | 16.82 ± 0.3 | 14.15 ± 0.621 |
| Lipid Content (mg/fly) | 0.02 ± 0.008 | 0.032 ± 0.002 | 0.055 ± 0.003 | 0.141 ± 0.006 | - |
| Glycogen Content (mg/fly) | 0.04 ± 0.002 | 0.053 ± 0.004 | 0.072 ± 0.005 | 0.157 ± 0.009 | - |
| Water Content (mg/fly) | 0.586 ± 0.01 | 0.761 ± 0.022 | 0.891 ± 0.01 | 1.076 ± 0.012 | - |
| Body Mass (mg/fly) | 0.796 ± 0.014 | 1.045 ± 0.027 | 1.263 ± 0.032 | 1.643 ± 0.035 | - |

Note – All data was collected at age 14 days from egg. All data represents phenotypic characterization of female flies from the 50 populations used in this study. Statistics for cardiac arrest rates were performed via Cochran-Mantel-Haenszel test. Statistics for remaining six parameters were calculated via linear mixed effects and estimated marginal means analysis. P-value significance cutoff of 0.05.

### 4.2 Desiccation Resistance, Glycogen Content and Water Content

Similar to the starvation resistance data, we found the same trend in desiccation resistance (Table 1, Fig 1C). For the A-type populations, the ACO1-5 and AO1-5 populations had an average desiccation survival time of 10.35 hours and 10.416 hours, respectively (Table S1). For the B-type populations. the B1-5 and BO1-5 populations had average desiccation survival times of 10.504 hours and 11.066 hours, respectively. The A-type and B-type populations did not significantly differ in desiccation resistance, and they collectively had the shortest desiccation survival times (p-value = 0.427, Table S2). The C-type populations, with their longer life-cycles, were observed to have increased desiccation resistance (14.67 survival hours), as reported in previous studies (e.g. Graves et al. 1992; Rose et al. 1992). The CO1-5 populations and nCO1-5 populations had average desiccation survival times of 14.709 and 14.631 hours, respectively. The C-type populations did have significantly higher survival times than the A-type and B-type populations on a pairwise basis, as determined using post hoc EMmeans analysis (p-values < 0.0001). In addition, when analyzing the three pairwise differences between long-established and recently derived populations within selection regimes, there are no statistically significant differences for desiccation resistance (p-values > 0.05, Table S3). For the T-type regime, the TDO1-5 populations and TSO1-5 populations had average desiccation survival times of 15.037 and 13.262 hours, respectively. Despite the 1.78 hour difference in survival time, this difference was not significant (p-value = 0.164). However, the 10 T-type populations do not differentiate from the 10 C-type populations (p-value = 0.376). Similar to the starvation resistance, the S-type populations clearly have the highest desiccation resistance, with average survival time of 16.83 hours (Table S1). In a desiccated environment, they survive longer than the next two more resistant groups, the C-type and T-type populations (p-values of 0.0007 and <0.0001, respectively, Table S2).

With glycogen and water contents being two of the main factors behind desiccation resistance (Fig 2), it is not surprising that we see similar trends as the desiccation resistance. The ACO and AO populations had similar average water contents of 0.572 mg/fly and 0.6 mg/fly (p-value = 0.202, Table S3). The B and BO populations also have similar average water contents of 0.776 mg/fly and 0.747 mg/fly (p-value = 0.548). The CO and nCO populations had similar average water contents of 0.919 mg/fly and 0.863 mg/fly (p-value = 0.114). The five SCO_A_ and five SCO_B_ populations had average water contents of 1.058 mg/fly and 1.094 mg/fly, respectively (p-value = 0.374). The S-type average water content was significantly higher than the average water content of the C-type populations (p-value = <0.0001). Regarding glycogen content, the relationships between the different regimes are the same as water content. The average glycogen contents for the ACO and AO populations were 0.035 mg/fly and 0.044 mg/fly, respectively (p-value = 0.1302). The average glycogen contents for the B and BO populations were 0.055 mg/fly and 0.050 mg/fly, respectively (p-value = 0.6081). The average glycogen contents for the CO and nCO populations were 0.068 mg/fly and 0.077 mg/fly, respectively (p-value = 0.367). The average glycogen contents for the SCO_A_ and SCO_B_ populations were 0.151 mg/fly and 0.161 mg/fly, respectively (p-value = 0.473). Just like the water content and desiccation resistance, the A-type population have the lowest glycogen content whereas the S-type populations clearly have the highest glycogen contents (Table 1; Table S1). Of all the between selection regime comparisons, only the A-type versus B-type glycogen content comparison was not significantly different (p-value = 0.0854). Unlike desiccation resistance, we did not measure glycogen content and water content for the T-type populations.

**Figure 2.**
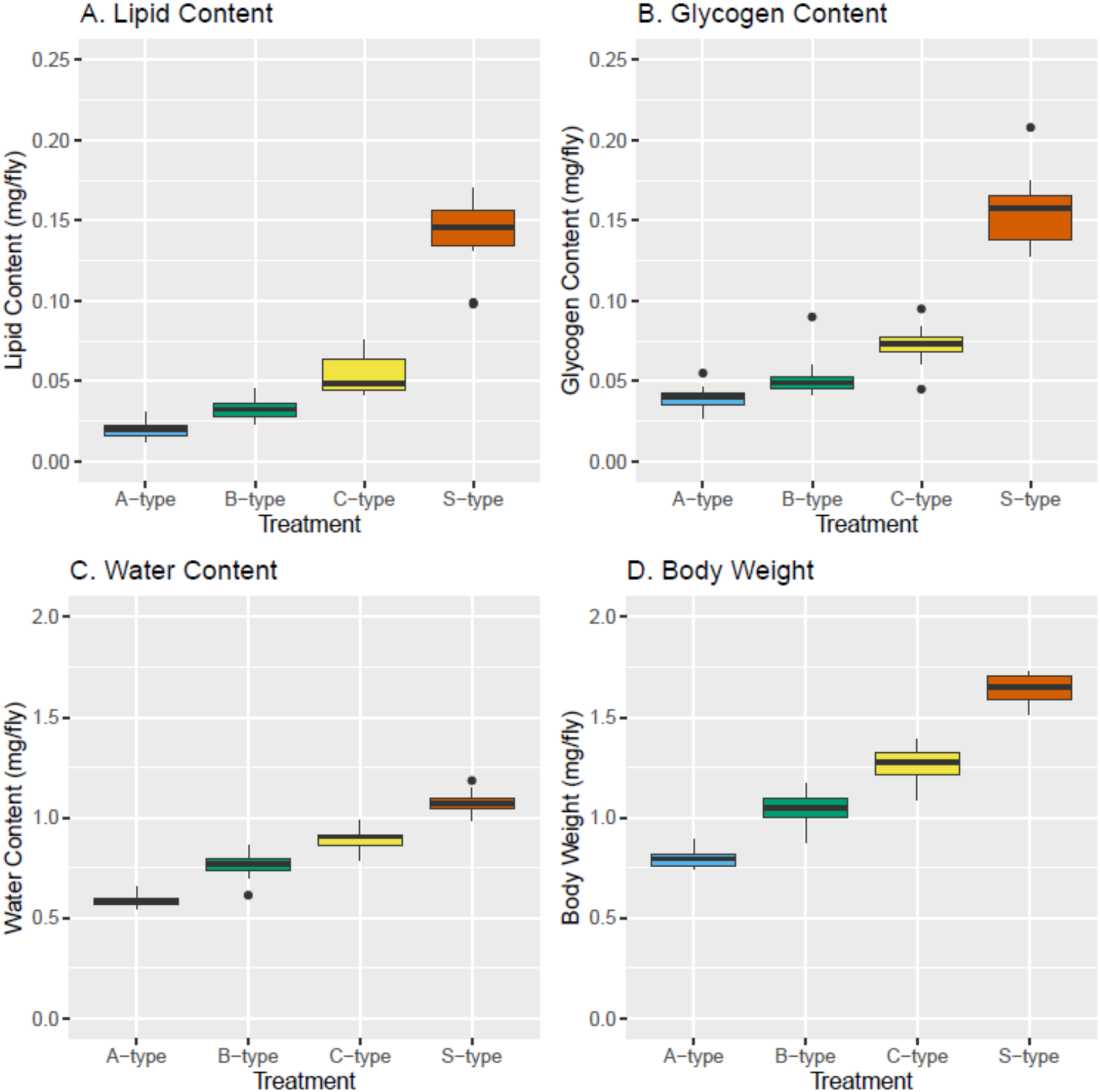
Summary boxplots of metabolic reserves and overall body weight for the four selection regimes. In lipid content (A), glycogen content (B), and water content (C), we see an upward trend across the selection regimes along generation cycle length. Additionally, selection for increased starvation resistance (S-type regime) has led to significantly increased metabolic reserves compared to their respective 28-day cycle length controls (C-type regime). Similarly, we observe the same trend and significances regarding the overall body weight of these flies, in which these three metabolic reserves entail majority of the body weight. (R-studio linear mixed effects model analysis).

### 4.3 Cardiac Arrest Rate

Populations with longer generation cycles were found to have lower cardiac arrest rates, with the exception of the S-type populations (Fig. 1A, Table 1). The A-type populations, which have the 10-day generation cycle, recorded the highest cardiac arrest rates. The five ACO populations had a 56.5% cardiac arrest rate, with the five AO populations had a 54% cardiac arrest rate (p-value = 0.474; Table S1 and S3). The second highest set of cardiac arrest rates were associated with the SCO-a (54% arrest rate) and SCO-b (52.7%) populations (p-value = 0.718). The B-type populations had intermediate cardiac arrest rates. The five B populations had an average cardiac arrest rate of 35.5% and the five BO populations had an average arrest rate of 39.5% (p-value = 0.224). The 10 T-type and 10 C-type populations have the lowest cardiac arrest rates. The five CO populations had an average arrest rate of 25.3%, whereas the five nCO populations had an average arrest rate of 27.3% (p-value = 0.518). The five TSO populations had an average arrest rate of 27.6%, whereas the five TDO populations had an average arrest rate of 25.8% (p-value = 0.557).

With no significant differences found within each of the five selection regimes (Table S3), we then compared the cardiac arrest rates between the five selection regimes (Table S2). There was no significant difference between the 10 C-type populations (average arrest rate of 26.33%) and the 10 T-type populations (average arrest rate of 26.7%, p-value = 0.895). The 10 S-type populations had an average cardiac arrest rate of 53.33%, and this was significantly higher than the B-type, T-type, and C-type average rates (p-value < 0.0001). The S-type cardiac arrest rate was not significantly different than the A-type cardiac arrest rate (p-value = 0.441). Lastly, the B-type populations had an intermediary average cardiac arrest rate of 37.5%, which was significantly lower than the A-type and S-type populations, and also significantly higher than the C-type and T-type populations.

### 4.4 SNP Identification and FLAM Analysis

The genomic analysis of the 40 populations (A-, B-, C-, and S-type populations) with the subsequent CMH analysis identified 1,258 50kb genomic windows with a representative SNP (Figure 3). The genomic analysis and subsequent CMH analysis of all 50 populations (now including T-type populations) identified 1,951 50kb genomic windows with a representative SNP (Figure 3). After identifying these representative SNPs between our populations, our FLAM analyses identified a total of 161 SNPs and further linked these SNPs with one of eight phenotypic parameters (Figure 4). The breakdown of which SNPs are potentially associated with the phenotypic parameters are broken down in Table S4. For example, the 50 population FLAM analysis with the cardiac pacing results yielded 18 SNPs. The 161 identified SNPs are located somewhat evenly across the five major chromosome arms (Figure 4, Table S5). Thirty-eight are located on chromosome arm 2L, 27 SNPs are located on chromosome 2R, 27 SNPs are located on chromosome 3L, 43 SNPs are located on chromosome 3R, and the remaining 26 SNPs are located on chromosome X. Checking for overlap between identified SNPs and known genetic regions via FlyBase genome browser, we found a total of 155 unique genes that fell within a ± 1 kilobase window around the SNP position (Table S6). Eighteen genes are associated with cardiac arrest rate results, The associated GO functions of these genes vary from cellular and organelle function to DNA and RNA expression elements (see Table S7).

**Figure 3.**
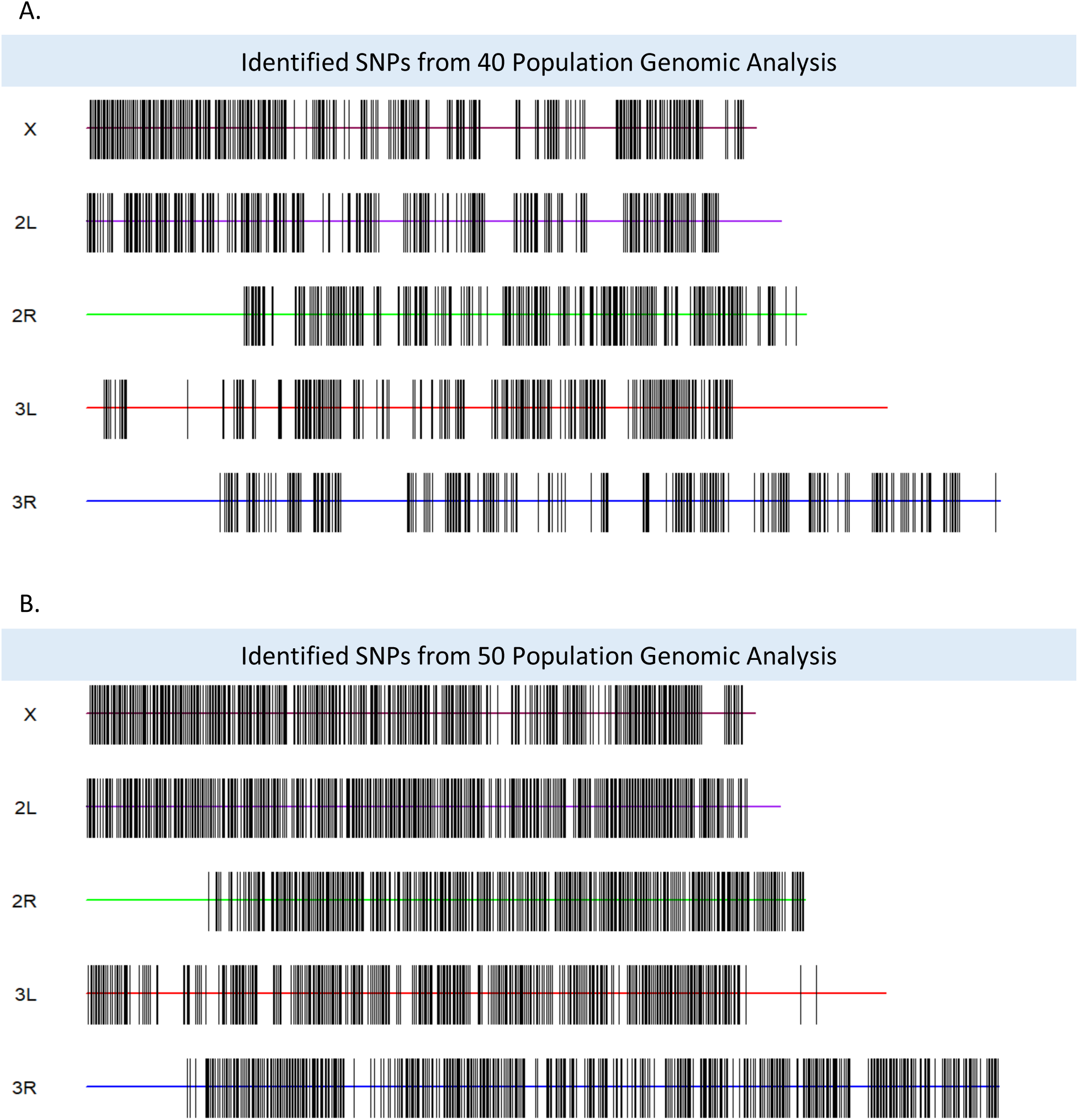
These genomic “barcodes” represent identified SNP sites across the five major chromosomal arms. (A) These chromosomal “barcode” images represents 1,258 SNP sites that were significantly different between the 40 populations (A-, B-, C-, and S-type populations) and fed into the FLAM analysis for the metabolic reserves and body weights. (B) These chromosomal “barcode” images represents 1,951 SNPs that were significantly different between the 50 populations (A-, B-, C-, S-, and T-type populations). Significantly differentiated SNPs between the populations are identified via Cochran-Mantel-Haenszel analysis.

**Figure 4.**
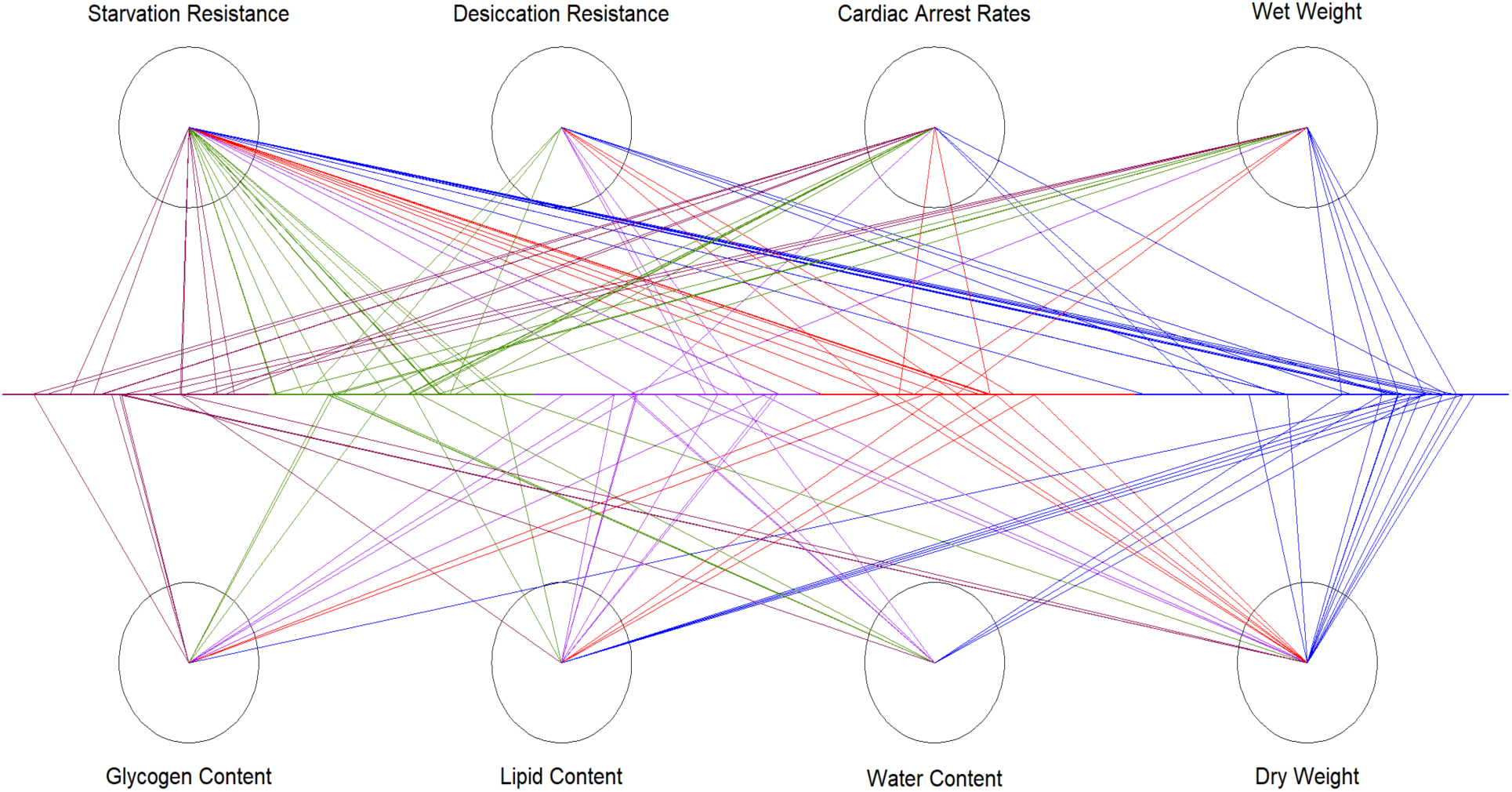
Mapping of FLAM-identified predictive SNPs to their respective physiological parameter. The five different colors represent the five different major chromosome arms (X: maroon; 2L: green; 2R: purple; 3L: red; 3R: blue). For each parameter, there are linked predictive SNPs from across the different chromosome arms. The predictability of one trait is not localized to one specific genomic region, yet alone, one specific chromosome arm.

## Discussion

### 5.1 Physiological Differentiation Among Selection Regimes

What we have observed in this study is corroboration of previous comparative and evolutionary physiology findings with regard to stress resistance, metabolic reserves, and life history, both those from our lab as well as studies from outside our lab (vid. Rose et al. 2004; cf. Gibbs and Gefen 2009). Our expectation is that the underlying biochemical mechanisms behind this evolutionary physiology of life-history are highly likely to be the same mechanisms as those found previously. For example, total stored calories and lipid content have been strongly correlated with starvation resistance (Djawdan et al. 1998), whereas trehalose and glycogen content are predictive for desiccation resistance and flight duration (Graves et al. 1992; Djawdan et al. 1998). In addition to carbohydrate content, decreasing water loss rates and increasing bulk water content are two mechanisms which evolve in response to selection for desiccation resistance (Gibbs et al. 1997; Archer et al. 2007). The A-type populations have been selected for accelerated development (10-day generation cycle); they age faster and are significantly smaller than both B-type and C-type populations (Chippindale et al. 2004). In our study, the A-type populations have the shortest survival times under starvation and desiccation, lowest metabolic reserves, and smallest body mass. The A-type populations also had the highest rate of pacing-induced cardiac arrests (55.25% arrests). The B-type populations, which have 14-day generation cycle, had intermediate phenotypic values in regards to the A-type, T-type and C-type populations. The T-type and C-type populations, which did not differ significantly from each other for the majority of the observed phenotypes, have 21-day and 28-day generation cycles, respectively. They have similar cardiac arrest rates, survival time during starvation, and survival time during desiccation. For example, the T-type and C-type populations have improved cardiac arrest rates (26.7% and 26.33% cardiac arrest rates, respectively) compared to the A-type (55.25%) and B-type (37.5%) populations. The 10 C-type populations had larger metabolic reserves and body masses than the A-type and B-type populations.

The correlations between the above mentioned physiological responses and generation cycle begin to breakdown in the S-type populations. These S-type populations were intensely selected for starvation resistance over a six-year span and have significantly larger amounts of metabolic reserves which explains the changes in body masses and survival times during starvation and desiccation. Unlike the other 40 populations, where we documented an inverse relationship between metabolic reserves and cardiac arrest rates, the S-type populations have the largest metabolic reserves and the second highest rate of cardiac arrests among these selection regimes (53.33% arrest rate).

So what underlying mechanisms could account for this spectrum of heart robustness? In an experiment looking at the effect of triglycerides, glycogen and water content on heart properties, it was determined that differences in the level of these macromolecules do not significantly affect cardiac robustness in populations with three or four-week life cycles (Shahrestani 2011). This suggests that body content differences are not as important as life-history differentiation in determining cardiac robustness. On a still larger scale than the study of Shahrestani, we show that longer-lived C-type populations have increased rates of heart recovery after pacing compared to shorter-lived A-type populations. However, there is only a certain extent to the lack of macromolecule impact on heart function. Regarding the breakdown of these aforementioned correlations in our S-type populations, we believe it is likely, but not proven, to be due to the substantial increased amount of lipids (0.38 mg of lipid higher than C-type populations). The impact of increased lipid concentration could affect cardiac function in various ways, either through increased hemolymph viscosity, changes in phospholipid homeostasis, or physical impingement of lipid stores on the cardiac tube (Hardy et al 2015; Hardy et al 2017; Kezos et al 2019).

Overall, we observed positive correlations between generation cycles, metabolic reserves, and survival times under stress. However, we also witnessed this correlation break down when examining heart robustness in populations selected to the extreme. As in Phelan et al. (2003), by characterizing multiple phenotypes for these 50 populations, we illustrate not just that there are costs that arise at high levels of stress resistance, but that the evolutionary relationship between two functional traits may be complex.

### 5.2 How Well Do Identified SNPs and Associated Genes Correlate with Physiological Function?

So what are the genes that might help explain the observed differences in physiological phenotypes? As mentioned, the identified SNPs by our FLAM analyses span the whole genome, with no specific region being responsible for a phenotypic trait (see Fig. 4, Table S5). Furthermore, one chromosome arm did not have significantly more causal SNPs than another. These results support the idea of a polygenic or network-like foundation underlying the phenotypic response to different selection pressures. For all of the phenotypes, with the exception of starvation resistance and dry weight, the identified SNPs for each phenotype were found somewhat evenly across the five major arms (Table S5). For starvation resistance, the chromosome arms 2L and 3R had 15 and 12 SNPs respectively, with the other arms having between four and eight SNPs. Similarly, for the dry weight associated SNPs, 11 of the SNPs were found on chromosome arm 3R.

Amongst the genes associated with identified SNPs, nearly a third were associated with multiple phenotypes (see Table S6). For instance, the gene alphabet (*alph*), which encodes a serine/threonine phosphatase involved in the regulation of the RAS/MAPK signaling pathway, is associated with both starvation resistance and dry weight phenotypes. Chromosome-associated protein D3 (*Cap-D3*) has previously been associated with chromatin binding activity and regulation of DNA metabolic processes. In our analyses, this gene and its associated SNP were identified as a causal determinant for glycogen content, water content, and overall wet mass data. Of the 30 genes with multiple associated phenotypes, 13 of those genes are associated with both water content and wet mass. Aside from tissue biomass, water content is the largest factor contributing to wet mass. In regards to genes associated with the cardiac arrest rate phenotype, only two were associated with another phenotype: starvation resistance (Dpr-interaction protein _K_ / *DIP-Kappa* and Glycerophosphate oxidase 3 / *Gpo3*).

Among the phenotypes, starvation resistance phenotype was associated with the greatest number of genes (47 genes). The previously identified functions of these genes include axongenesis, DNA/RNA binding and repair, embryonic development, morphogenesis, phospholipid transportation, and neuropeptide activity, to list a few examples (see Table S7). As we generally see with all of the phenotypes, the genes associated with their differentiation have had a wide array of functions attributed to them, functions not always transparently linked to the associated phenotype. The gene *Dop1R1* is a dopamine receptor that is associated with responses to starvation and sucrose. *Mondo* is a transcription factor that is involved in lipid and carbohydrate metabolism, lipogenesis, and both glucose and triglyceride homeostasis. The gene *wunen* (wun) is a lipid phosphate phosphatase that is required for germ cell migration and survival, tracheal septate junction function and heart formation. It is also associated with KEGG (Kyoto Encyclopedia of Genes and Genomes) pathways such as glycerolipid metabolism, glycerolipid metabolism, ether lipid metabolism, and sphingolipid metabolism (Table S7). There are two enriched biological process GO terms linked with these starvation resistant-associated genes, response to sucrose and female germ-line stem cell asymmetric division. Sucrose is a major component of Drosophila metabolism and a source of energy. It has been shown that high-sucrose diets in flies can disrupt metabolic homeostasis and promote metabolic disorders such as insulin resistance, obesity and diabetes (Musselman et al 2011; Musselman and Kühnlein 2018; Musselman et al. 2019). Additionally, the identified genes *dally*, *ds*, and *cyclin E* are associated with the hippo signaling pathway, which according to the gProfiler analysis (Raudvere et al 2019), is significantly enriched (p-value 0.0037; see Table S7).

And if we look for enriched GO terms and pathways within the identified lipid-associated genes via DAVID bioinformatics, we see enrichment for various GO terms and protein domains such as BMP signaling, cGMP biosynthesis, cGMP-mediated signaling guanylate cyclases (Wang et al 2011, Morrell et al 2016). The pathways highlighted here, along with the Wnt and Hippo signaling enrichment in the starvation resistance genes, are thought to play a role in cardiogenesis and function (Vogler and Bodmer 2015; Zhang and Del Re 2017; Foulquier et al. 2018).

And when we focus on the set of genes associated with the cardiac arrest data, only one KEGG pathway term was significantly enriched, glycerophospholipid metabolism (genes CG7311 and *rdgA*; p-value = 0.022). Glycerophospholipids are components within lipid bilayers of cellular components, and altering the metabolism of these lipids could have effects on membrane fluidity, protein function, ion channel activity, signal transduction and secondary messaging. Although not significant, small GTPase mediated signal transduction, a GO term with implications for cardiac development and function, was nearly enriched (p-value = 0.072). Two of the genes associated with the enrichment of this term are *sponge* and CG42541, with the latter known to enable GTP binding activity and calcium channel regulatory activity.

Genes associated with desiccation, glycogen, and water content led to only a few enriched GO terms and/or KEGG pathways. Three of the desiccation resistant-associated genes (CG4936, *Eip75B*, *InaC*) led to a significant enrichment for zinc ion binding (p-value = 0.014). *InaC* and *Zasp52* are also associated with calcium-dependent activities and alpha-actinin binding, two components that are linked to *Drosophila* heart function. In regards to glycogen content, there were a few cell junction and cellular component GO terms that were significantly enriched. However, of more interest are the enriched KEGG pathways. Similar to cardiac arrest rates, we once again see glycerophospholipid metabolism, in addition to glycerolipid metabolism, ether lipid metabolism and sphingolipid metabolism. Lastly, one biological process GO term was nearly significantly enriched within the water content gene set, negative regulation of Notch signaling pathway (p-value = 0.051). Notch signaling is a highly conserved pathway that is very important for cell-to-cell communication. It is necessary for proper cell differentiation and proliferation during both development and adult life. Dysregulation of Notch signaling has been associated with disorders such as leukemia, multiple sclerosis, Alagille syndrome, cardiovascular disorders and many others (Lasky and Wu, 2005; Shang et al 2016; Christopoulos et al 2021).

### 5.3 *Accuracy of FLAM-derived Predictive Models for Physiological Traits (*Thomas, I need you to fill in the details missing, such as X and Y number of populations)

After identifying significant SNPs for each physiological trait with our FLAM analysis (Figure 4, Table S4), we were interested in testing the accuracy of the generated predictive models. For each of our physiological traits, the data was separated into two parts, a training set including 40 populations (8 from each regime) and a testing set including the remaining 10 populations (2 from each regime). Splitting populations into the training set or testing set was repeated for different combinations to ensure each population was included at least once in the testing set. For the physiological traits in which we only had data from the 40 populations, the training set used 32 populations and the testing set included the remaining 8 populations. Using our training sets of populations, we include phenotypic data as outcomes and the SNP data for the focal predictors as inputs. With each specific fit, we used SNP data from the test set populations to predict their phenotypes and then compared those predicted values to the observed phenotypic values. The correlation values between the predicted phenotype and observed phenotype values for each of the eight physiological traits were determined. We found strong correlations between the predicted values and observed for all of the traits. All but one of the correlation values are higher than 0.9, with the correlation value for the desiccation resistance trait being 0.84. It should be noted that desiccation resistance was the least differentiated phenotype in the current study; so its lower genomic predictability makes intuitive sense. In any case, these high predictive correlations show that FLAM allows us to determine which differentiated genomic regions are chiefly contributing to specific types of phenotypic differentiation.

Previous studies in our lab (Mueller et al 2018; Barter et al. unpublished) suggested that high levels of replication are necessary to detect most causal loci using FLAM analysis. These articles used 20 populations, and although they detected causal loci, they suggested that the number of populations that should be used in FLAM analyses should exceed 40 populations. The present physiological study is the first to utilize the genomics of up to 50 populations, depending on the physiological trait, in FLAM analyses. By including these additional populations, we have increased our power to detect causal loci, as shown by the higher predictive-model correlation values reported here as compared to our lab’s previous work. Understanding the genomic complexity of adaptation is slowly becoming achievable as we improve our machine learning tools (e.g. FLAM), increase replication at the population level, and incorporate multi-omic data sets (i.e. genomics, transcriptomics, proteomics and metabolomics).

### 5.4 Understanding the Molecular Genetics underlying Evolutionary Physiology

In Barter et al. (unpublished), our lab used the three levels of genomic, transcriptomic, and phenotypic data for 20 experimentally-evolved *Drosophila* populations (the here-described 10 A-type and 10 C-type populations). Using FLAM to investigate how these three levels of data are interconnected with one another opens opportunities to address long-standing and unanswered questions about the molecular foundations of functional and physiological adaptation. Our lab is knocking on the door of determining whether phenotypic adaptation follows a simple one-to-one relationship between gene and phenotype or is more polygenic or even network-like.

Mueller et al (2018), Barter et al (unpublished), and our study show that multiple loci across the genome underlie the differentiation of physiological characters in response to experimental evolution. In other words, a single gene is not solely or even primarily responsible for the evolution of individual physiological traits. Thus physiological genetics outside of the limited arena of large-effect mutants should be expanded to allow for polygenic networks of causation. Barter et al (unpub) report that single differentiated genomic regions affected the patterns of transcription of multiple genes, while conversely each transcript’s expression level was predicted by numerous genomic regions. Their results and ours are in conformity with Wright’s (1980) vision of complex genetic networks underlying adaptations.

While it is certainly not going to be the only machine learning tool of value, FLAM evidently allows the effective parsing of the molecular genetic basis of physiology. The present study establishes the complexity of the interactions between the genome and functional physiology. It is doubtful that evolutionary physiology in the genomic era can continue to make progress in the face of this complexity without some type of machine learning to forge connections bridging the genome and physiology.

## Acknowledgments

We thank James Hicks for helpful discussions and comments on the experiment. We also thank Michael Mulligan for use of laboratory equipment. We are extremely grateful to the many undergraduate research students who contributed to the stock maintenance, experimental assays, and data collection as this experiment occurred over five years.

## Supplemental Tables and Figures

**Table S1.** Phenotypic Characterization of ACO, AO, B, BO, CO, nCO, SCOA, SCOB, TSO and TDO Populations

| Population | Cardiac Arrest<br>Rate | Survival during<br>Starvation (h) | Survival during<br>Desiccation (h) | Lipid Content<br>(mg/fly) | Glycogen Content<br>(mg/fly) | Water Content<br>(mg/fly) | Body Mass<br>(mg/fly) |
| --- | --- | --- | --- | --- | --- | --- | --- |
| ACO1 | 0.55 | 31.348 | 9.547 | 0.020 | 0.035 | 0.578 | 0.785 |
| ACO2 | 0.53 | 33.253 | 11.525 | 0.031 | 0.034 | 0.591 | 0.806 |
| ACO3 | 0.63 | 34.132 | 9.585 | 0.021 | 0.040 | 0.548 | 0.753 |
| ACO4 | 0.60 | 43.319 | 10.079 | 0.024 | 0.027 | 0.543 | 0.745 |
| ACO5 | 0.53 | 28.419 | 11.017 | 0.023 | 0.040 | 0.600 | 0.823 |
| AO1 | 0.50 | 38.673 | 9.851 | 0.014 | 0.046 | 0.601 | 0.819 |
| AO2 | 0.63 | 30.762 | 9.965 | 0.015 | 0.040 | 0.563 | 0.755 |
| AO3 | 0.55 | 46.730 | 11.371 | 0.020 | 0.044 | 0.611 | 0.822 |
| AO4 | 0.58 | 21.387 | 10.663 | 0.012 | 0.036 | 0.572 | 0.769 |
| AO5 | 0.45 | 33.399 | 10.230 | 0.019 | 0.055 | 0.654 | 0.889 |
| B1 | 0.33 | 31.927 | 11.149 | 0.037 | 0.051 | 0.804 | 1.103 |
| B2 | 0.35 | 48.407 | 11.225 | 0.030 | 0.045 | 0.703 | 0.962 |
| B3 | 0.38 | 50.978 | 11.376 | 0.024 | 0.049 | 0.832 | 1.131 |
| B4 | 0.38 | 48.660 | 9.735 | 0.027 | 0.042 | 0.765 | 1.043 |
| B5 | 0.35 | 40.683 | 9.033 | 0.035 | 0.090 | 0.774 | 1.060 |
| BO1 | 0.40 | 45.309 | 10.772 | 0.035 | 0.049 | 0.781 | 1.091 |
| BO2 | 0.45 | 55.350 | 11.074 | 0.045 | 0.053 | 0.614 | 0.877 |
| BO3 | 0.40 | 50.461 | 10.809 | 0.029 | 0.060 | 0.739 | 1.001 |
| BO4 | 0.43 | 47.154 | 11.338 | 0.036 | 0.042 | 0.738 | 1.014 |
| BO5 | 0.30 | 47.374 | 11.338 | 0.023 | 0.048 | 0.863 | 1.170 |
| CO1 | 0.3 | 64.533 | 14.233 | 0.064 | 0.061 | 0.918 | 1.327 |
| CO2 | 0.217 | 72 | 13.31 | 0.068 | 0.076 | 0.91 | 1.308 |
| CO3 | 0.283 | 62.133 | 15.7 | 0.052 | 0.084 | 0.984 | 1.387 |
| CO4 | 0.2 | 68.966 | 14.633 | 0.044 | 0.045 | 0.871 | 1.22 |
| CO5 | 0.267 | 66.8 | 15.667 | 0.063 | 0.075 | 0.912 | 1.295 |
| nCO1 | 0.233 | 68 | 13.517 | 0.042 | 0.07 | 0.933 | 1.327 |
| nCO2 | 0.25 | 61.517 | 12.069 | 0.076 | 0.072 | 0.83 | 1.206 |
| nCO3 | 0.367 | 63.733 | 15.467 | 0.045 | 0.095 | 0.789 | 1.092 |
| nCO4 | 0.25 | 56.667 | 16.167 | 0.045 | 0.078 | 0.862 | 1.212 |
| nCO5 | 0.267 | 57.067 | 15.933 | 0.044 | 0.068 | 0.903 | 1.257 |
| SCO1A | 0.533 | 159.286 | 15.867 | 0.146 | 0.128 | 1.059 | 1.699 |
| SCO2A | 0.5 | 185.286 | 16.6 | 0.157 | 0.159 | 0.983 | 1.533 |
| SCO3A | 0.5 | 175.6 | 16.667 | 0.142 | 0.144 | 1.016 | 1.516 |
| SCO4A | 0.55 | 129.704 | 18.333 | 0.098 | 0.167 | 1.150 | 1.711 |
| SCO5A | 0.617 | 105.067 | 18.967 | 0.145 | 0.157 | 1.081 | 1.654 |
| SCO1B | 0.4 | 138.667 | 15.567 | 0.17 | 0.128 | 1.084 | 1.73 |
| SCO2B | 0.617 | 186.591 | 16.733 | 0.132 | 0.208 | 1.05 | 1.574 |
| SCO3B | 0.55 | 171.333 | 14.7 | 0.155 | 0.136 | 1.102 | 1.651 |
| SCO4B | 0.583 | 144.207 | 17.5 | 0.099 | 0.175 | 1.185 | 1.721 |
| SCO5B | 0.483 | 120.966 | 17.367 | 0.158 | 0.16 | 1.048 | 1.645 |
| TSO1 | 0.278 | 72.400 | 11.667 | - | - | - | - |
| TSO2 | 0.233 | 82.222 | 12.345 | - | - | - | - |
| TSO3 | 0.278 | 73.857 | 16.833 | - | - | - | - |
| TSO4 | 0.303 | 66.207 | 12.367 | - | - | - | - |
| TSO5 | 0.289 | 73.000 | 13.100 | - | - | - | - |
| TDO1 | 0.278 | 70.533 | 13.138 | - | - | - | - |
| TDO2 | 0.233 | 73.379 | 14.700 | - | - | - | - |
| TDO3 | 0.256 | 66.667 | 15.467 | - | - | - | - |
| TDO4 | 0.256 | 67.867 | 14.467 | - | - | - | - |
| TDO5 | 0.267 | 66.800 | 17.414 | - | - | - | - |

Note – All data was collected at age 14 days from egg. All data represents phenotypic characterization of female flies from the 50 populations used in this study.

**Table S2.** P-Value Summary of Physiological Comparisons Between Selection Regimes

| Regime Comparison | Cardiac Arrest Rate | Survival during Starvation (h) | Survival during Desiccation (h) | Lipid Content (mg/fly) | Glycogen Content (mg/fly) | Water Content (mg/fly) | Body Mass (mg/fly) |
| --- | --- | --- | --- | --- | --- | --- | --- |
| A v B | <0.0001 | 0.056 | 0.4274 | 0.0614 | 0.0854 | <0.0001 | <0.0001 |
| A v C | <0.0001 | <0.0001 | <0.0001 | <0.0001 | 0.0001 | <0.0001 | <0.0001 |
| A v T | <0.0001 | <0.0001 | <0.0001 | - | - | - | - |
| A v S | 0.441 | <0.0001 | <0.0001 | <0.0001 | <0.0001 | <0.0001 | <0.0001 |
| B v C | <0.0001 | 0.0075 | <0.0001 | 0.0017 | 0.009 | <0.0001 | <0.0001 |
| B v T | <0.0001 | 0.0003 | <0.0001 | - | - | - | - |
| B v S | <0.0001 | <0.0001 | <0.0001 | <0.0001 | <0.0001 | <0.0001 | <0.0001 |
| C v T | 0.895 | 0.2532 | 0.3763 | - | - | - | - |
| C v S | <0.0001 | <0.0001 | 0.0007 | <0.0001 | <0.0001 | <0.0001 | <0.0001 |
| S v T | <0.0001 | <0.0001 | <0.0001 | - | - | - | - |

Note – All data was collected at age 14 days from egg. All data represents phenotypic characterization of female flies from the 50 populations used in this study. Statistics for cardiac arrest rates were performed via Cochran-Mantel-Haenszel test. Statistics for remaining six parameters were calculated via linear mixed effects and estimated marginal means analysis. P-value significance cutoff of 0.05.

**Table S3.** P-Value Summary of Physiological Comparisons Within Selection Regimes

| Regime Comparison | Cardiac Arrest Rate | Survival during Starvation (h) | Survival during Desiccation (h) | Lipid Content (mg/fly) | Glycogen Content (mg/fly) | Water Content (mg/fly) | Body Mass (mg/fly) |
| --- | --- | --- | --- | --- | --- | --- | --- |
| ACO v AO | 0.474 | 0.9289 | 0.8961 | 0.0194 | 0.1302 | 0.2016 | 0.333 |
| B v BO | 0.224 | 0.2386 | 0.2832 | 0.5371 | 0.6081 | 0.5478 | 0.621 |
| CO v NCO | 0.518 | 0.0809 | 0.9353 | 0.3371 | 0.3666 | 0.1144 | 0.0949 |
| TSO v TDO | 0.557 | 0.1517 | 0.1642 | - | - | - | - |
| SCOA v SCOb | 0.718 | 0.9501 | 0.2779 | 0.7537 | 0.473 | 0.3736 | 0.4266 |

Note – All data was collected at age 14 days from egg. All data represents phenotypic characterization of female flies from the 50 populations used in this study. Statistics for cardiac arrest rates were calculated via Cochran-Mantel-Haenszel analysis. Statistics for remaining six parameters were calculated via linear mixed effects analysis. P-value significance cutoff of 0.05.

**Table S4.** FLAM-identified SNPs

| Phenotype | Fly Gene (Gene Symbol + Flybase Gene Number) |
| --- | --- |
| Cardiac Arrest Rates | 2L: 12716836, 2L: 12838690, 2L: 13379865, 2L: 13708673, 2L: 17171614, 2L: 9613765, 2R: 9750392, 3L: 15755566, 3L: 7366021, 3R: 28857479, 3R: 6267037, 3R: 9177571, X: 19116519, X: 20418434, X: 2574202, X: 3917297, X: 8931345, X: 9051130 |
| Starvation Resistance | 2L: 10852190, 2L: 13708673, 2L: 13801704, 2L: 15737489, 2L: 15840679, 2L: 15850477, 2L: 16363018, 2L: 18994810, 2L: 21289452, 2L: 21316786, 2L: 21966012, 2L: 3250503, 2L: 6057588, 2L: 686080, 2L: 754853, 2R: 18104121, 2R: 22630990, 2R: 22659312, 2R: 9402359, 3L: 11673750, 3L: 14801223, 3L: 15679948, 3L: 15755566, 3L: 15901865, 3L: 271302, 3L: 5706848, 3L: 8848734, 3R: 14136565, 3R: 14173934, 3R: 24218319, 3R: 24272460, 3R: 24533483, 3R: 25014947, 3R: 27254873, 3R: 27778389, 3R: 27814935, 3R: 28594243, 3R: 29592239, 3R: 970361, X: 16400146, X: 16503180, X: 19809899, X: 21391673, X: 6249098, X: 6250430, X: 8376500 |
| Lipid Content | 2L: 13299384, 2L: 21364394, 2R: 16918965, 2R: 21980126, 2R: 22377354, 2R: 7348693, 2R: 9346339, 2R: 9497227, 3L: 11058790, 3L: 16267836, 3L: 19787725, 3L: 19801671, 3R: 25014947, 3R: 26796210, 3R: 28354792, 3R: 30251512, X: 16478317 |
| Desiccation Resistance | 2L: 16714276, 2L: 3322632, 2L: 6899490, 2R: 15801805, 2R: 16901488, 2R: 20320038, 3L: 12897123, 3L: 17993912, 3L: 5554959, 3R: 13464317, 3R: 20032686, 3R: 23520205 |
| Glycogen Content | 2L: 10872456, 2L: 5534334, 2L: 5984306, 2R: 21252994, 2R: 2594243, 2R: 7348693, 2R: 7375173, 2R: 9402359, 3L: 5606038, 3L: 9573889, 3R: 26735338, X: 10061948, X: 10795391, X: 10958705, X: 2904284 |
| Water Content | 2L:13967269, 2L:5469295, 2L:5534334, 2L:5984306, 2R:11143168, 2R:18552526, 2R:8701989, 2R:8953809, 3R:19029858, 3R:22768596, 3R:26649206, X:13469718 |
| Body Mass (Wet) | 2L: 143491, 2L: 15424664, 2L: 5534334, 2L: 5984306, 2R: 8701989, 3L: 12548691, 3L: 8685631, 3R: 19029858, 3R: 22768596, 3R: 24277382, 3R: 26649206, 3R: 26651324, 3R: 29594182, X: 11037787, X: 11073347, X: 13469718, X: 16552013 |
| Dry Mass | 2L: 21595427, 2R: 12947802, 2R: 15051195, 2R: 21040098, 3L: 11058790, 3L: 12615746, 3L: 16267836, 3L: 19801671, 3L: 5706848, 3L: 7029070, 3R: 10513324, 3R: 14054031, 3R: 22768596, 3R: 24224131, 3R: 24277382, 3R: 25371699, 3R: 26344475, 3R: 28354792, 3R: 29594182, 3R: 30212315, 3R: 31254345, X: 10795391, X: 11113145, X: 16503180 |

Note – Genes were identified by comparing overlap between the FLAM-identified SNP position and *Drosophila* genome browser available on Flybase (www.flybase.org)

**Table S5.**
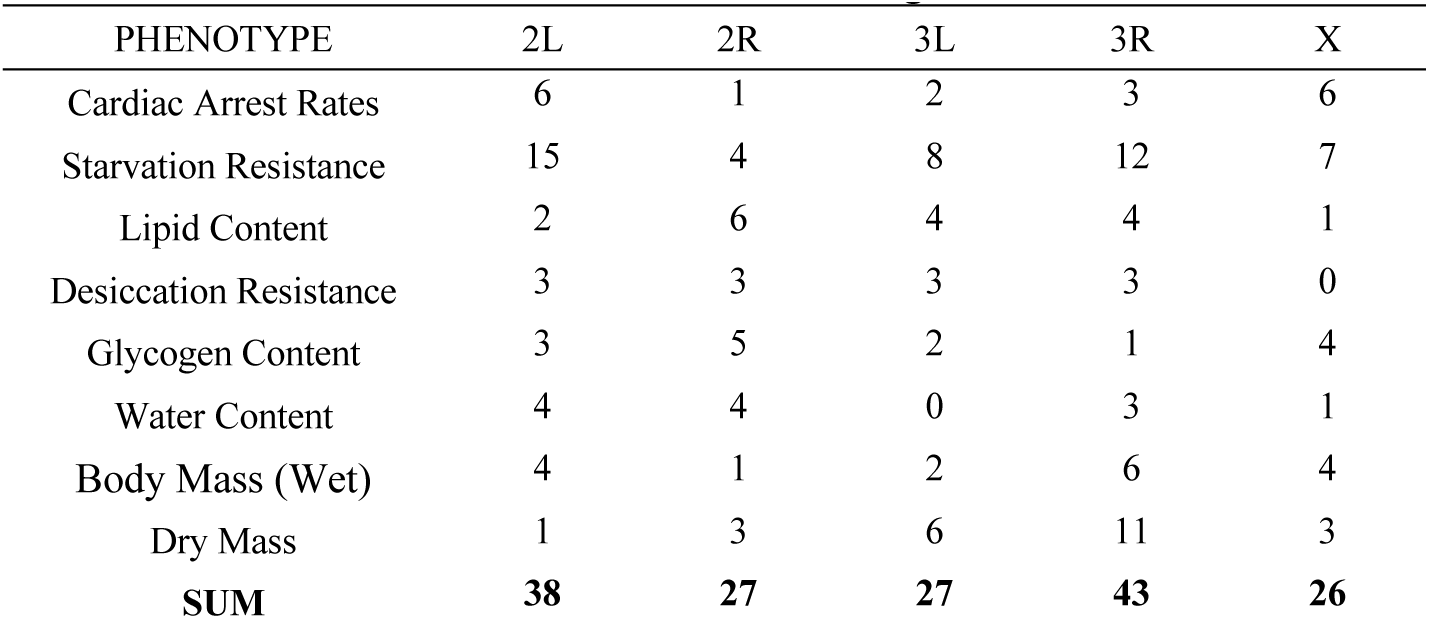
FLAM-identified Gene Distribution Among Chromosomal Arms

**Table S6.**
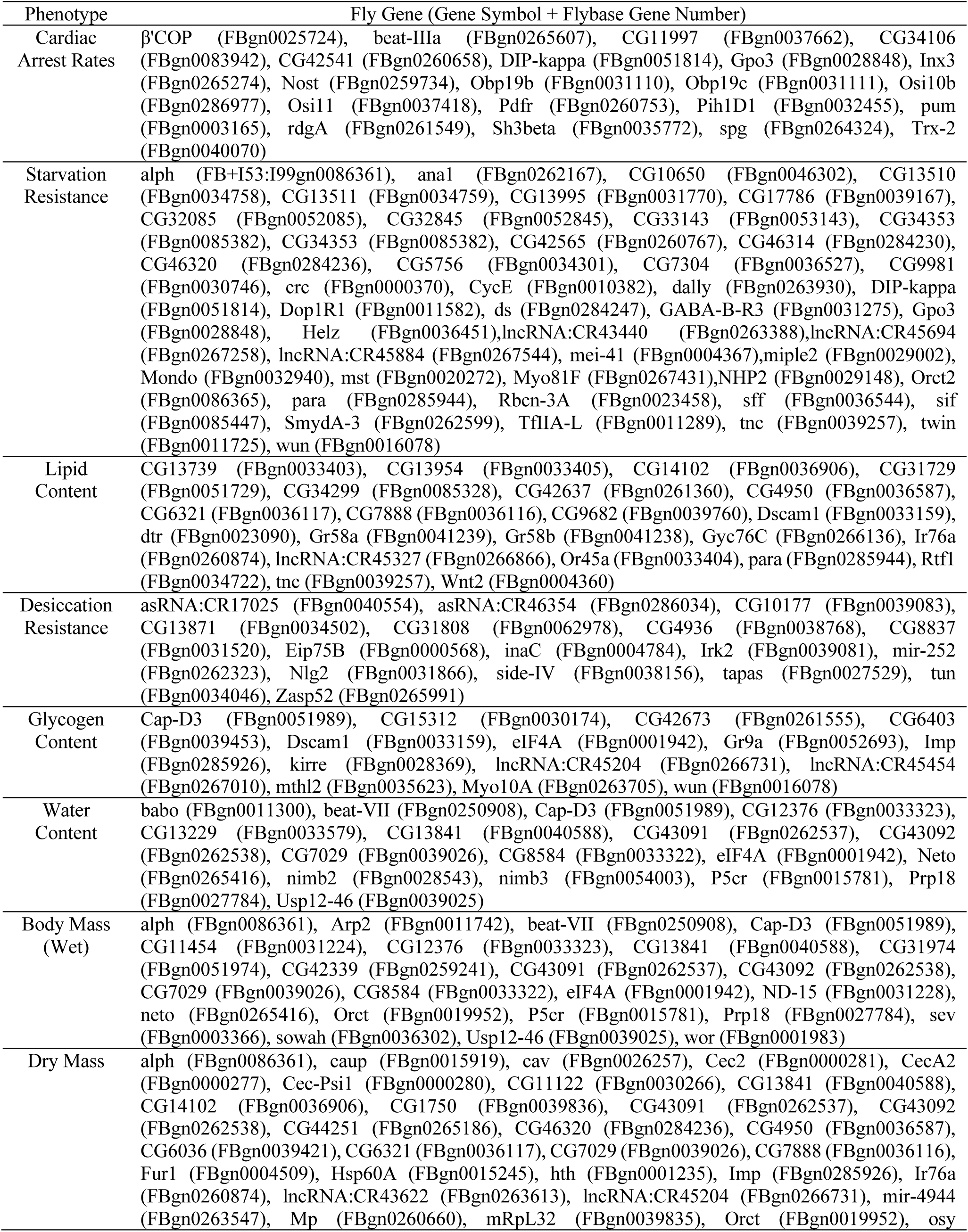
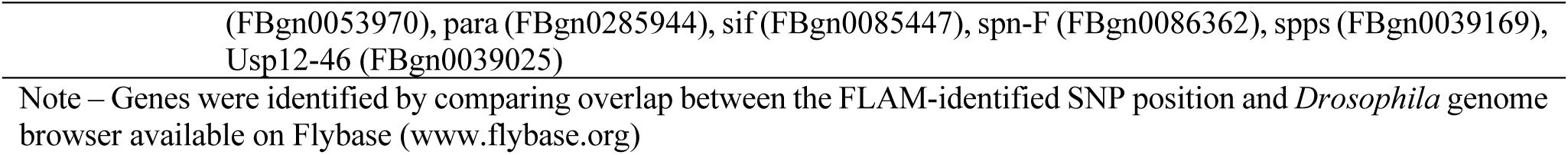
Identified Genes Associated With FLAM-Identified SNPs

**Table S7.**
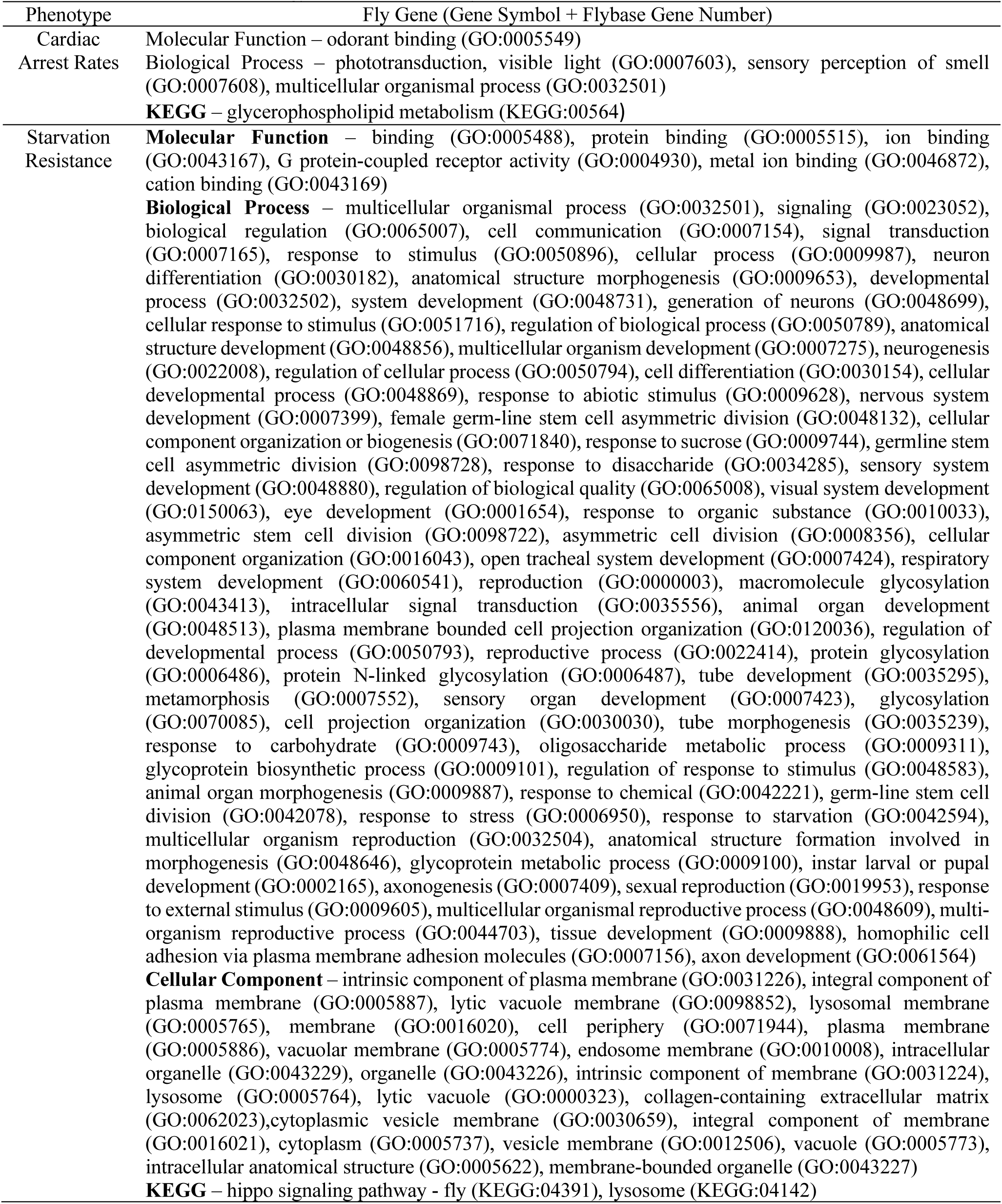
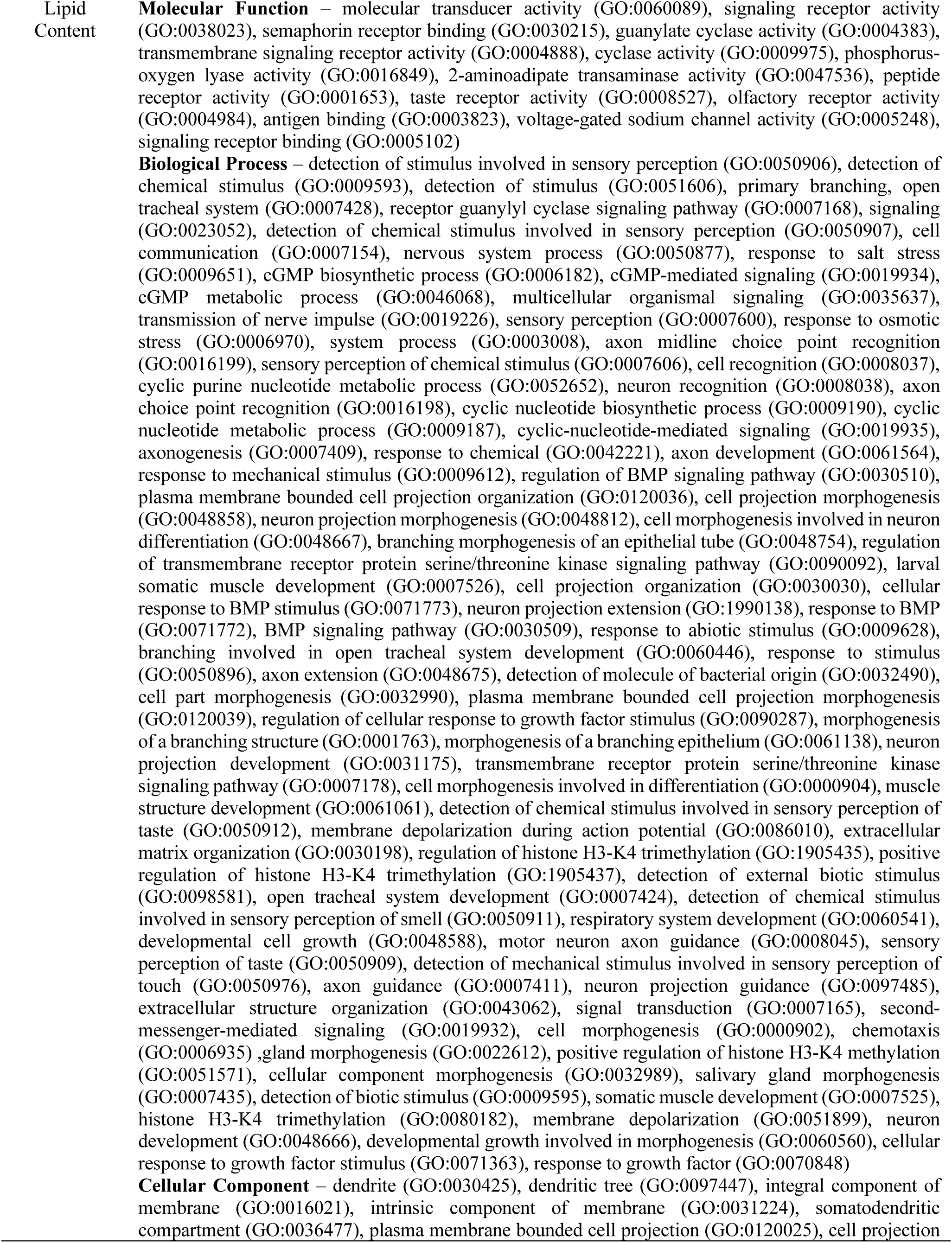
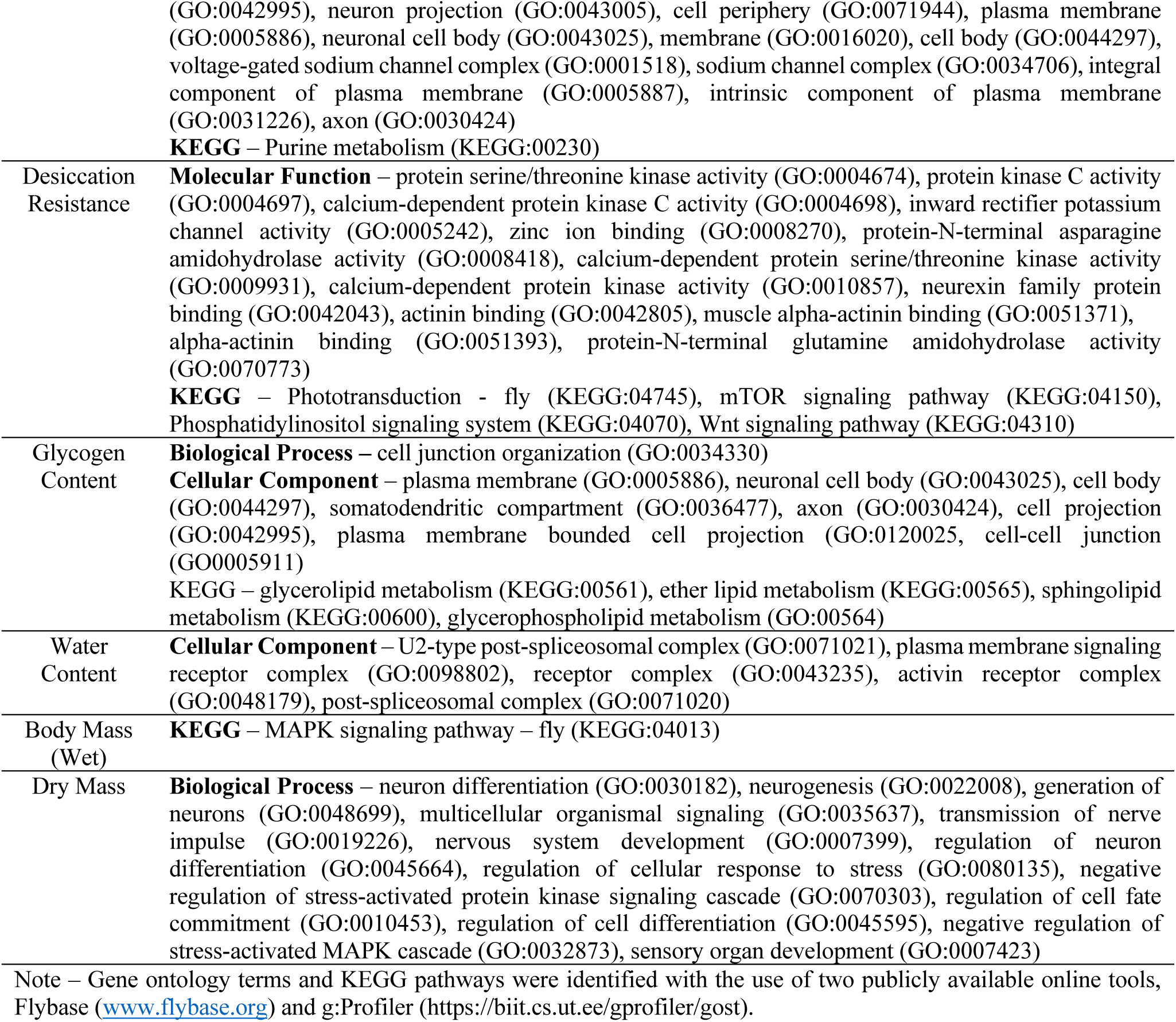
Gene Ontology and KEGG Pathway Associated with FLAM-identified Genes

